# Secondhand homes: Woodpecker cavity location and structure influences secondary nester’s success

**DOI:** 10.1101/2020.09.11.293860

**Authors:** Faith O. Hardin, Samantha Leivers, Jacquelyn K. Grace, David M. Cairns, Tyler Campbell, Brian Pierce, Michael L. Morrison

**Author notes:** **Author contributions:** FH conceived, designed, and analyzed the project. SL, JG, DC, and BP provided insights on the analysis and interpretation throughout the process. All authors contributed towards editing. Corresponding Author: Faith Olivia Hardin.

## Abstract

1. Understanding how ecosystem engineers influence other organisms has long been a goal of ecologists. Woodpeckers select nesting sites with high food availability and will excavate and then abandon multiple cavities through their lifetime. These cavities are crucial to secondary cavity nesting birds (SCB) that are otherwise limited by the availability of naturally occurring cavities.
2. Our study examined the role food resources have on the nest site location and home range size of woodpeckers, and the respective influence woodpeckers and the construction of cavities have on the nesting success of SCB.
3. Using five years of avian point count data to locate golden-fronted woodpeckers (GFWO: *Melanerpes aurifrons*), we correlated insect availability with GFWO home range size and determined differences in insect availability between GFWO occupied and unoccupied sites, while recording nesting success (success: ≥ 1 fledgling) for the GFWO and common SCB in south Texas: Black-crested Titmouse (*Baeolophus atricristatus*), Ash-throated Flycatcher (*Myiarchus cinerascens*), Brown-crested Flycatcher (*Myiarchus tyrannulus*), and Bewick’s Wren (*Thryomanes bewickii*). We used model averaging to fit species-specific logistic regression models to predict nest success based on cavity metrics across all species.
4. Sites occupied by GFWO had a higher biomass of insects in orders Coleoptera, Hymenoptera, and Orthoptera than unoccupied sites, and there was a negative correlation between the availability of these insect orders and home-range size. GFWO had increased nest success in trees with increased vegetation cover and lower levels of decay, while SCB had higher levels of nesting success in abandoned GFWO cavities opposed to naturally occurring ones, and in trees with low decay.
5. Our results suggest that SCB may be drawn to nest in abandoned woodpecker cavities where they have higher rates of nest success compared to natural cavities. Additionally, the prevalence for GFWO to excavate cavities in trees with lower levels of decay contradicts previous literature and may indicate a novel temperature trade-off, with live trees requiring more energy to excavate, but providing more protection from high breeding season temperatures in arid and semi-arid areas.

## 1. INTRODUCTION

Ecosystem engineers control the availability of resources for other species by causing physical state changes in biotic or abiotic materials (Jones, Lawton & Shachak, 1994; Wright, Jones, & Flecker, 2002; Buse et al., 2008). Given the important role they play in local environments, the literature surrounding ecosystem engineers is historically focused on how their actions affect other species (Jones et al., 1994; Robles & Martin, 2013; Tarbill, Manley, & White, 2015; Wiebe, 2017), but little research has been done concerning external factors that influence the engineers themselves (see Mikusinski, 2006; Jusino, Lindner, Banik, & Walters, 2015). Importantly, little has been done to investigate how ecosystem engineers choose breeding and young rearing grounds (Nilsson, Johnsson, & Tjernberg, 1991; Garmendia, Cárcamo, & Schwendtner, 2006). Understanding these driving factors is essential to understanding the ecology of not only the ecosystem engineers themselves, but the organisms that rely on them for their own breeding and nesting grounds as well.

The modifications made by ecosystem engineers have far-reaching consequences and directly impact not only ecological associations, but also the behavior of animals within an ecosystem. For example, animal movement and community composition may be altered by the actions of local ecosystem engineers (Lill & Marquis, 2003; Bangert & Slobodchikoff, 2004). In this way, ecosystem engineers can indirectly influence local trophic levels through multi-level environmental modifications, such as by influencing local invertebrate diversity and abundance, which in turn may increase foraging opportunities for other vertebrates (Lill & Marquis, 2003; Bangert & Slobodchikoff, 2004), or by providing more suitable species specific habitat for nesting (Showalter & Whitmore, 2002)

Although insects themselves can act as ecosystem engineers (Bell & Whitmore, 1997; Lill & Marquis, 2003; Bangert & Slobodchikoff, 2004), they can also act as crucial resources for other ecosystem engineers at higher trophic levels (Hess & James, 1998; Pechacek & Kristin, 2004). For example, declines in insect richness and abundance have been reported with parallel declines in a number of insectivorous ecosystem engineers, such as woodpeckers (Lister & Garcia, 2018, Møller, 2019, Karr, 1976; Benton, Bryant, Cole, & Crick, 2002; Rioux Paquette, Pelletier, Garant & Bélisle, 2014; Narango, Tallamy, & Marra, 2017; Bowler, Heldbjerg, Fox, Jong, & Böhning-Gaese, 2019). Therefore, ecosystem engineering activities may be better understood by looking at the distribution and abundance of their food resources.

Woodpeckers are avian ecosystem engineers that have a large proportion of insects in their diet (Jones et al., 1994; Tarbill et al., 2015), and control the location, construction, and availability of nesting cavities, a limiting resource for secondary cavity nesting birds (SCB; i.e. species that require a cavity to nest in but cannot create the cavity themselves). Woodpeckers are primary excavators of nesting cavities, often creating multiple cavities within their home range per year to avoid predation, external parasite buildup, and cavity wood degradation (Loye & Carroll 1998; Husak & Husak, 2002; Wiebe, 2017). Once abandoned, these cavities are used by a variety of secondary cavity nesting species (Martin & Eadie, 1999, Pakkala, Tiainen, Piha, & Kouki, 2019). Woodpeckers select nesting sites based on characteristics that protect their eggs and nestlings from predation, tending to nest high in moderately to heavily decayed trees with wide diameters at breast height (DBH), and with limited vegetation covering the cavity entrance (vegetation cover, Mannan, Meslow, & Wight, 1980; Li & Martin, 1991; Loye & Carroll, 1998; Newlon, 2005; Jusino et al., 2016). Additionally, the shape of woodpecker cavities functions to exclude nest predators by having small entrance holes and deep depths (Sedgwick & Knopf, 1990; Li and Martin, 1991; Martin, Aitken, & Wiebe, 2004; Rhodes, O’donnell, & Jamieson, 2009). Given the nest construction preferences of woodpeckers, the cavities they leave behind are often superior nesting spaces when compared to naturally occurring cavities, both of which are used by SCB (Martin & Li, 1992; Maziarz, Broughton, & Wesolowski, 2017).

Woodpecker resources can be defined both in terms of food (mainly wood burrowing insects, largely in the order Coleoptera) and in the number of trees suitable for excavation (Bonnot, Millspaugh, & Rumble, 2009; Rota, Rumble, Lehman, Kesler, & Millspaugh, 2015). These resources have been shown to be directly linked to woodpecker nest site location and home range sizes (e.g. the area used by a bird in its daily movements) (Worton, 1989; Powell, 2000; Wiktander, Olsson, & Nilsson, 2001; Pasinelli, 2007). For example, the Black-backed woodpecker (*Picoides arcticus*) selects nesting sites based on infestations of the mountain pine beetles (*Dendroctonus ponderosae*) (Rota et al., 2015), and the Three-toed woodpecker’s (*Picoides dorsalis*) home range size is negatively correlated with the number of trees with suitable DBH for cavity excavation (Pechacek & d’Oleire-Oltmanns, 2004). However, no studies to date have looked at the impact of food resources on both the nest site location and home range sizes of woodpeckers, which in turn directly impacts neighboring SCB.

The Golden-fronted woodpecker (GFWO, *Melanerpes aurifrons*), is a poorly studied, medium sized bird, whose range extends from Central America to Texas (Wetmore, 1948; Sauer, Link, Failon, Pardieck, & Ziolkowski, 2013; Schroeder, Boal, & Glasscock, 2013). GFWO numbers are in decline across their Texas distribution, and are considered a species of concern in the Texas Wildlife Action Plan (Bender, 2007). As with other woodpecker species, GFWO act as ecosystem engineers, providing nesting cavities for SCB throughout their range (Husak & Maxwell, 1998). Determining the factors that influence the nest site location and construction of cavities is crucial to not only understand the conservation needs of GFWO, but also for the conservation and basic ecology of SCB that may rely on the cavities GFWO create.

To investigate relationships between the GFWO and local SCB nesting successes, we conducted an observational study on GFWO nesting success (≥ 1 fledgling) in relation to nesting site locations, home range sizes, local insect biomass, and cavity construction, along with the nesting success of the four most common SCB in our study area, the Black-crested Titmouse (BCTI; *Baeolophus atricristatus*), Ash-throated Flycatcher (ATFL; *Myiarchus cinerascens*), Brown-crested Flycatcher (BCFL; *Myiarchus tyrannulus*), and Bewick’s Wren (BEWR; *Thryomanes bewickii*) in the southern Texas Tamaulipan Brushlands (Baumgardt, Morrison, Brennan, Pierce, & Campbell, 2019).

The objectives of our study were to determine 1) the role of insect availability in nest site location and home range size of GFWO, 2) the role of nest metrics (e.g. DBH, vegetation cover) in the nesting success of GFWO and the four species of SCB, and 3) if SCB tended to nest more in abandoned woodpecker cavities and had differing nesting success in abandoned woodpecker cavities compared to natural cavities. We predicted that 1) insect abundance would be greater at GFWO occupied sites versus GFWO unoccupied sites and that home range size would be negatively correlated with the availability of insect orders commonly eaten by birds, 2) the same cavity metrics would influence nest success in both GFWO and SCB species and 3) that SCB would tend to nest in, and have higher nest success in abandoned woodpecker cavities compared to natural cavities, and that abandoned woodpecker cavities would share characteristics making them more suitable for nesting birds, compared to natural cavities.

## 2. MATERIALS AND METHODS

### 2.1 Study Area

Our study was conducted on the East Foundation’s ~61,000 ha San Antonio Viejo (SAV) ranch located in Jim Hogg and Starr counties, ~25 km south of Hebbronville, south Texas. This area is representative of the Tamaulipan/Mezquital Thornscrub ecological region containing unique plants and animal communities within brush covered dunes, grasslands punctuated with clusters of trees, and open woods of mesquite (*Prosopsis glandulosa*). Annual rainfall during the study year (2019) for this region was ~30 cm and the mean temperature during the breeding season (March - July) was ~27.8° C (PRISM Climate Group 2019), similar to the 30 year norm for this region (PRISM Climate Group 2019). The SAV supports approximately 70 residential bird species and 45 migratory species (Baumgardt et al., 2019).

### 2.2 Nest Location and Monitoring

We used the East Foundation’s extensive long-term breeding bird dataset, constructed over 6 years, to create a heat map of areas most likely to contain nesting GFWO (Baumgardt et al., 2019). We then used the Point Density tool in ArcGIS version 10.3 (Environmental Systems Research Institute, Redlands, CA, USA) to take a 500 m^2^ fishnet sample, and interpolate density values across our study location. Within areas of high GFWO density, we placed 12 1-km^2^ survey plots (Figure S1) and from mid-April to late May, 2019 we visited each plot four times using the spot mapping technique to locate nesting GFWO (Martin & Geupel, 1993).

After locating GFWO nests, we searched 150 m^2^ grids centered around each nest every 3-5 days between April and July 2019 to document active SCB nests (Rodewald, 2004). To select GFWO unoccupied sites, we placed 150m^2^ grids 300 m away from occupied sites that had the same vegetation association but no observed GFWO activity (sightings, calling, drilling, foraging, and nesting) and searched for SCB nests in the same way. The vegetation associations were determined by the East Foundation’s hierarchical vegetation classification system, created in 2011-2012 where a vegetation association was defined by the dominant and subdominant species (Snelgrove, Dube, Skow & Engeling, 2013). To determine SCB nesting tendencies and any differences in cavity metrics between abandoned woodpecker cavities and natural cavities, we recorded and monitored all empty cavities we found in each grid throughout the breeding season.

We monitored each SCB and GFWO nest every 2-5 days to determine nest success; a nest was considered successful if ≥1 fledgling was observed outside the nest. After fledging, we measured the following nest metrics that have historically been predictors of cavity nesting success: the height of the nest measured from the center of the cavity opening to the base of the tree (height), the tree’s DBH, diameter of the cavity opening (opening), the depth of the cavity (depth), and decay ranking (decay), where a rank of one indicated a live tree and rank seven indicated a dead tree with no branches, bark, and soft stem (Dobkin, Pretare, & Pyle, 1995; Bonar, 2001; Cockle, Martin, & Wesolowski, 2011; Berl. Edwards, & Bolsinger, 2015). Because increased vegetation cover may be detrimental for cavity nesting birds (Schaaf, 2020), we used 0.5 x 0.5 m^2^ cover boards to estimate the percentage of vegetation cover at each cavity (Nudds, 1997; Chotprasertkoon, Pierce, Savini, Round, Sankamethawee, & Gale, 2017).

### 2.3 Insect Sampling and Home range delineation

To determine if GFWO were choosing nesting sites and home range sizes based on available insects, we compared home range sizes to the available insect biomass within. Home range size was estimated by constructing minimum convex polygons (MCPs) on a randomly chosen subset of the home ranges (n = 24). We constructed MCPs by recording male movements over four, 30-minute visits that began after observing a male leave their nest (Dudley & Saab, 2007). We recorded 120 observation points for each male and built MCPs using the minimum bounding geometry tool in ArcGIS version 10.3 (Environmental Systems Research Institute, Redlands, CA, USA)

Within the same subset of home ranges, along with the associated unoccupied sites, we quantified the availability of insects with an array of 11 sweep net sampling locations from the center of the site (0 m) outwards in 15 m increments to 150 m (see Figure S2), visiting each site once per week from May to mid-July 2019 (Doxon, Davis, & Fuhlendorf, 2011). We sorted the insects by order, dried them using an Elite Eliminator Heater set at 55°C, and weighed them every 24 hours until their mass stabilized.

### 2.4 Statistical analysis

#### 2.4.1. Insect availability

We averaged insect mass over the seven visits across sampling locations within a home range and summed all sampling locations per site to get a single measure of insect order biomass per site. We used Mann-Whitney U t-tests to determine differences (P = 0.05) in insect abundance between sites occupied by GFWO and unoccupied sites, and used Spearman’s Rho to test for significant correlations between each insect order’s biomass and each male GFWO’s home range size (Field et al., 2012).

#### 2.4.2. GFWO Nest Success

We created logistic regression models in RStudio version 1.15.2, (R Core Team 2013) with the package *car* (Fox & Weisberg, 2019) using recorded cavity metrics to predict GFWO nest success. We considered variance inflation factors (VIFs) >5 as indicators of multicollinearity between variables and z-scaled all continuous variables to account for varying units of measurement (O’Brien, 2007). To create candidate models, we used the *MuMIn* package (Barton, 2020) in R to generate a model selection table (Burnham & Anderson, 2002; Field, Miles, & Field, 2012), and evaluated model fit using AIC adjusted for small sample sizes (AICc) (Burnham & Anderson, 2002). Models that had ≥10% of the weight of the top model were considered candidate models for model averaging (Burnham & Anderson, 2004; Mazerolle, 2006). Using the R package *AICcmodavg* (Mazerolle, 2020) we estimated the parameter coefficients through model averaging and determined which parameters were significant using P ≤ 0.05 and corresponding confidence intervals.

#### 2.4.3 SCB Nest Success

To compare the structure of abandoned woodpecker cavities to natural cavities we used Welch’s tests for each set of measurements taken on all cavities encountered (Field et al., 2012). We then followed the same steps to create species specific logistic regression and model averages for the four SCB (Nemes, Jonasson, Genell, & Steineck, 2009; Field et al., 2012). Observations on the ATFL and the BCFL were combined given the similarity of their body metrics and life history traits, and hereafter are referred to as ATBC (Cardiff and Dittmann 2000). We used the same six cavity metrics, with the addition of whether the nest was located in an abandoned woodpecker cavity or a natural cavity (cavity type). As before, we used the R packages *MuMIn* and *AICcmodavg* to evaluate candidate models and average parameter coefficients per species.

## 3. RESULTS

### 3.1 Insects define GFWO localities

We collectively spent 560 hours recording GFWO activities and found 55 GFWO nests, along with an additional 2,880 observation hours to define GFWO home ranges. We spent 220 hours collecting insect samples across 24 of these home ranges and 24 unoccupied equivalent ranges, and found that insect orders Coleoptera (W = 19, P < 0.001), Orthoptera (W = 13, P < 0.001), and Hymenoptera (W = 186, P < 0.036) had significantly higher masses on GFWO occupied sites than unoccupied sites. All other insect orders were not significantly different.

GFWO home range sizes were negatively correlated with the same three orders of insects, Coleoptera (P < 0.001, rho = −0.74, n = 24), Orthoptera (P = 0.007, rho = −0.55, n = 24), and Hymenoptera (P = 0.009, rho = −0.53, n = 24) (see Figure 1). The biomass of Phasmatodea was positively correlated (P = 0.045, rho = 0.41, n = 24) with GFWO home range size, and all other insect orders were not significantly correlated.

**Figure 1:**
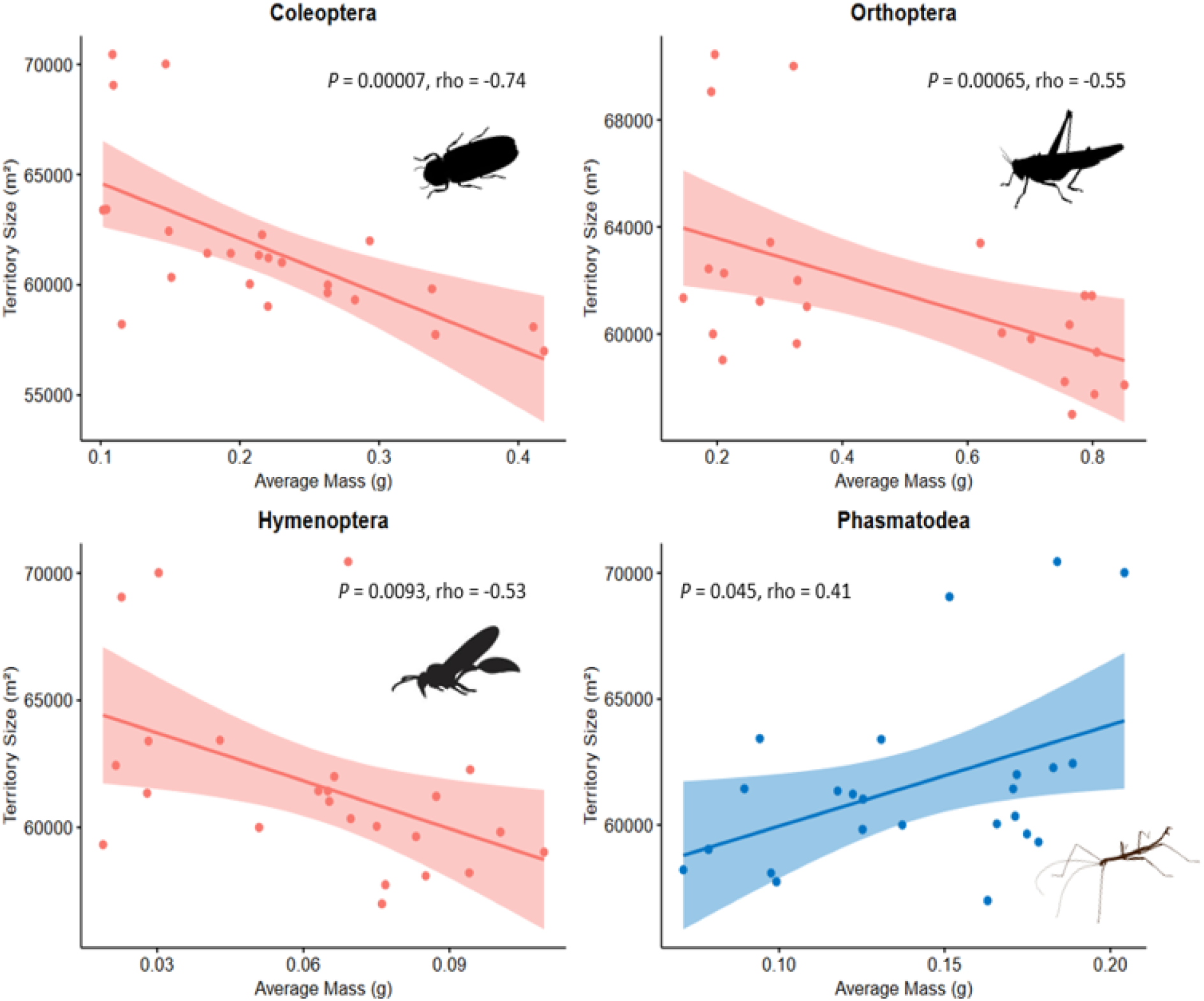
Scatter plots of Golden-fronted woodpecker home range size (m^2^) correlated with average mass (g) of significant insect orders. Shaded areas represent 95% confidence intervals. Data collected with sweep nets on the San Antonio Viejo Ranch, East Foundation in south Texas, during the summer of 2019.

### 3.2 GFWO nest success

The mean height for a GFWO cavity within our study was 2.3 m ± 0.26, the mean DBH of the nesting tree was 52 cm ± 6.2, the mean cavity diameter was 9 cm ± 0.8, the mean depth was 7 cm ± 0.7, and the mean vegetation cover was 43% ± 6.3. Over 25% of GFWO nests were in trees with decay class 1 (Table 1).

**Table 1:** Nesting tree decay (1 = live tree, 7 = dead, decayed tree), for each cavity nesting bird found within the study. Count and percent of that species within each decay rank are shown for each species of secondary cavity nesting bird, along with the primary cavity nesting bird, the Golden-fronted woodpecker. The data on the Ash-throated and Brown-crested Flycatchers were combined due to similar life history traits between species. Data was collected on the San Antonio Viejo Ranch, East Foundation in south Texas during the summer of 2019.

| Species | Decay |  |  |  |  |  |  |
| --- | --- | --- | --- | --- | --- | --- | --- |
|  | 1<br>(%) | 2<br>(%) | 3<br>(%) | 4<br>(%) | 5<br>(%) | 6<br>(%) | 7<br>(%) |
| Ash-throated/Brown-crested Flycatcher | 14<br>(13.7) | 11<br>(10.8) | 16<br>(15.7) | 23<br>(22.5) | 19<br>(18.6) | 16<br>(15.7) | 3<br>(2.9) |
| Black-crested Titmouse | 7<br>(17.9) | 5<br>(12.8) | 5<br>(12.8) | 3<br>(7.7) | 10<br>(25.6) | 6<br>(15.4) | 3<br>(7.7) |
| Bewick's Wren | 16<br>(20.3) | 10<br>(12.7) | 13<br>(16.5) | 15<br>(19) | 14<br>(17.8) | 11<br>(13.9) | 0<br>(0) |
| Golden-fronted Woodpecker | 14<br>(25.5) | 8<br>(14.5) | 7<br>(12.7) | 4<br>(7.3) | 7<br>(12.7) | 9<br>(16.4) | 6<br>(10.9) |

No VIFs were >5, thus all predictors were entered into the global model (see Table S1 for candidate model selection). Model averaging suggested that GFWO nests were less likely to be successful as decay increased (β = −0.91), and were more likely to be successful as vegetation cover increased (β = 0.10) (Table 2). Looking at the magnitude of effect, decay was ten times stronger at predicting successful nests for GFWO than vegetation cover, though both were significant. Notably, with every unit increase in decay (ranked 1-7) nest success for the GFWO dropped 0.41.

**Table 2:** Model averaged estimates with 95% confidence intervals (CI) for variables retained in the candidate model sets that predicted cavity nesting bird nesting success. All continuous variables used to create candidate models were z-scaled. Decay was ranked 1 = live tree, 7 = dead, decayed tree. Cavity Type = whether the nest was located in an abandoned woodpecker cavity or a naturally occurring one, DBH = diameter of the nesting tree at breast height. Flycatchers = combined observations of Ash-throated and Brown-crested flycatchers. Data was collected on the San Antonio Viejo Ranch, East Foundation in south Texas during the summer of 2019. Bootstrapping was used to obtain CI. SE is standard error and bolded variables are significant (P < 0.05)

|  |  |  |  | 95% CI |  |
| --- | --- | --- | --- | --- | --- |
| | Model<br>averaged $\beta$ | SE | P | Lower | Upper |
| Golden-fronted woodpecker (n = 55) |  |  |  |  |  |
| Decay | -0.91 | 0.41 | 0.015 | -1.71 | -0.1 |
| Vegetation Cover | 0.09 | 0.05 | 0.028 | -0.001 | 0.19 |
| DBH | 0.12 | 0.3 | 0.362 | -0.48 | 0.71 |
| Diameter of Opening | 0.05 | 0.33 | 0.445 | -0.59 | 0.69 |
| Height | 0.02 | 0.28 | 0.472 | -0.52 | 0.56 |
| Depth | 0.02 | 0.18 | 0.46 | -0.33 | 0.37 |
| Bewick's wren (n = 79) |  |  |  |  |  |
| Decay | -0.03 | 0.14 | 0.421 | -0.30 | 0.24 |
| Vegetation Cover | 0.06 | 0.02 | 0.002 | 0.02 | 0.10 |
| DBH | 0.63 | 0.49 | 0.104 | -0.34 | 1.59 |
| Diameter of Opening | -0.04 | 0.18 | 0.408 | -0.40 | 0.31 |
| Height | 0.01 | 0.17 | 0.480 | -0.34 | 0.33 |
| Depth | < 0.01 | 0.17 | 0.500 | -0.34 | 0.34 |
| Cavity Type (natural) | 1.92 | 0.95 | 0.023 | 0.05 | 3.78 |
| Flycatchers (n = 102) |  |  |  |  |  |
| Decay | -0.40 | 0.19 | 0.018 | -0.77 | -0.03 |
| Vegetation Cover | < 0.01 | 0.01 | 0.383 | -0.01 | 0.01 |
| DBH | < 0.01 | 0.14 | 0.498 | -0.27 | 0.27 |
| Diameter of Opening | -0.63 | 0.39 | 0.056 | -1.40 | 0.14 |
| Height | 0.06 | 0.19 | 0.385 | -0.32 | 0.43 |
| Depth | -0.05 | 0.17 | 0.388 | -0.39 | 0.29 |
| Cavity Type (natural) | 3.54 | 0.77 | < 0.001 | 2.02 | 5.05 |
| Black-crested titmouse (n = 39) |  |  |  |  |  |
| Decay | -1.02 | 0.41 | 0.008 | -1.83 | -0.21 |
| Vegetation Cover | 0.03 | 0.03 | 0.180 | -0.03 | 0.08 |
| DBH | 0.07 | 0.29 | 0.403 | -0.49 | 0.63 |
| Diameter of Opening | 0.02 | 0.21 | 0.460 | -0.39 | 0.43 |
| Height | -0.05 | 0.29 | 0.429 | -0.63 | 0.52 |
| Depth | < 0.01 | 0.21 | 0.497 | -0.42 | 0.42 |
| Cavity Type (natural) | 2.53 | 1.28 | 0.025 | 0.03 | 5.04 |

Note: Candidate models were chosen if they had an AICc weight $\geq 10\%$ of the AICc weight of the top model.

### 3.3 Cavities and SCB nesting success

Across all cavities found, whether a nest had been initiated in it or not, abandoned woodpecker cavities were significantly different than natural cavities: abandoned woodpecker cavities were built 42% higher in less decayed trees with 20% larger DBH than natural cavities and had 18% higher vegetation cover (Table 3). The size of the entrance hole and the depth of the cavity were not significantly different between nest types.

**Table 3:** Results of Welch’s t-test comparing differences between abandoned woodpecker cavities (AWC) and natural cavities (NC). DBH = diameter of the nesting tree at breast height, Decay (1 = live tree, 7 = dead, decayed tree). Data was collected on the SAV Ranch, East Foundation during 2019.

|  | P | t | AWC |  | NC |  |
| --- | --- | --- | --- | --- | --- | --- |
|  |  |  | Average | (±) | Average | (±) |
| **Decay** | **< 0.001** | **9.3** | **3** | **0.3** | **4** | **0.3** |
| **Vegetation Cover (%)** | **< 0.001** | **6.4** | **50** | **1.6** | **41** | **1.8** |
| **DBH (cm)** | **< 0.001** | **8.3** | **63.1** | **1.5** | **50.2** | **1.2** |
| Opening (cm) | 0.321 | 20.1 | 13.6 | 4.2 | 15.2 | 6.7 |
| **Height (m)** | **< 0.001** | **22.1** | **1.9** | **0.2** | **1.1** | **0.15** |
| Depth (cm) | 0.297 | 9.7 | 20.2 | 5.7 | 18.4 | 7.3 |

Model averaging for the BEWR suggested that cavity type was 15 times stronger at predicting successful nests than vegetation cover, though both were significant (Table 2; see Table S1 for candidate model selection), with nests more likely to be successful as vegetation cover increased (β = 0.06), and if nests were built in an abandoned woodpecker cavity over a natural cavity (β = 0.95). Model averaging for both the BCTI and the ATBC suggested that decay and the cavity type were significant predictors for nest success. As with the GFWO, with every unit increase in decay, nest success dropped 0.19 for ATBC and 0.41 for BCTI. Again, cavity type was the strongest predictor; cavity type was 3 times stronger at predicting nest success than decay for the BCTI, and was 4 times stronger than decay for the ATBC. Across SCB species, cavity type was the strongest predictor of nest success.

## 4. DISCUSSION

Decades of field observations in a range of bird species suggest the importance of insects to birds during the breeding season, as protein demands are increased while producing eggs and provisioning nestlings (Capinera, 2011, Vitz & Rodewald, 2012). We identify correlations between food resources and GFWO nest site location and home range size, along with nest cavity characteristics that facilitate successful broods and reveal the importance of abandoned woodpecker cavities for secondary cavity nesting birds. Additionally, our results suggest a novel trade-off between excavating live trees versus dead/decaying trees, evident in the differences in nest success between natural cavities and abandoned woodpecker cavities.

### Resource driven site location

All recorded orders of insects collected within our study were found at all occupied and unoccupied site types, though not every insect order was found at each sweep netting location, nor at every visit. Previous literature has indicated that Coleoptera and Hymenoptera are in high proportions of woodpecker diets (Beckwith & Bull, 1985; Hess & James, 1998; Fayt, Machmer, & Steeger, 2005; Pechacek & Kristin, 2010), and as we predicted in our first objective, the biomass of both of these insect orders were higher around GFWO nests than unoccupied sites and increases in their biomass corresponded with decreased GFWO home ranges, up to 15,000 m^2^. In addition, we found similar relationships between Orthoptera and GFWO sites and home ranges.

Our findings indicate that resource availability (e.g. insect biomass) may be driving the location and home range sizes of this ecosystem engineer, as GFWO nests were located in areas that corresponded with insect availability, and home ranges shrank in correlation with increases in those same insect orders. This is in accordance with previous literature which indicates that woodpeckers reduce their defended areas when resources were abundant, and chose nesting sites based on resource availability (Pasinelli, 2000; Tingley, Wilkerson, Bond, Howell, & Siegel., 2014). The differences we found in insect biomass between occupied and unoccupied sites were most likely due to fine scale variation in vegetation and water availability indistinguishable by our vegetation associations (Huang, Zhao, & von Gadow, 2015).

### Interconnected nesting success

In our second and third objectives, we predicted that the same cavity metrics that influenced GFWO nest success would also influence SCB, and that SCB would have higher nest success in abandoned woodpecker cavities. As predicted, all SCB had higher nest success rates in abandoned woodpecker cavities than in natural cavities and cavity type was the strongest predictor for all species, with the BEWR having the least impact, followed by the BCTI, and largest influence on ATBC. Additionally, GFWO had higher success in trees with lower decay and higher vegetation cover, which was mirrored in SCB; BCTI and ATBC were more likely to produce fledglings in trees with low decay, and BEWR were more likely to produce fledglings in cavities with high vegetation cover. The BEWR was the only species not impacted by decay, potentially explained by its generalistic nesting behavior (Taylor, 2003). We observed successful BEWR nests built in metal pipes or direct sun, thus experiencing wide temperature swings throughout the day, indicating that unstable nesting environments may be a deterrent for other cavity nesting birds, but not this species.

Also in line with our third objective, we predicted that abandoned woodpecker cavities would share characteristics making them better nesting cavities than natural ones. To this, SCB within our study had higher success rates within abandoned woodpecker cavities (81-93%), than in natural cavities (41-56%). The structure of abandoned woodpecker cavities present on our sites were distinctly different from their natural counterparts; on average they were significantly higher in trees, of lower decay, smaller DBH, and increased vegetation cover, all characteristics that protect eggs and fledglings from shifting internal temperatures and predation (Copeyon, 1990; Ojeda, Suarez, & Kitzberger, 2007; Pakkala et al., 2019). Considering that SCB are reliant on pre-existing cavities to create their nests, the factors that drive the creation and design of woodpecker cavities may then dictate the success of local SCB.

### Tree decay and vegetation cover: a possible role for temperature

We found a higher than expected number of GFWO nests within live trees. Previous literature on woodpecker nesting ecology has indicated a preference for excavating cavities in partially to fully decayed trees, which require less energy and time than dense, live wood (Conner, Miller, & Adkisson, 1976; Cockle et al., 2011; Blanc & Martin, 2012). However, these studies have focused on temperate regions such as northwestern, northeastern United States, Canada, and European countries where breeding season temperature rarely exceeds 35° C and occasionally reach freezing during the early spring (Conner et al., 1976; Blanc & Martin, 2012; Seavy, Burnett, & Taille, 2012). In contrast, the mean breeding season temperature at our study site in southern Texas was 27.8° C and daytime temperatures frequently reached over 42.2° C Currently, there is little information on how cavity nesting birds regulate nest temperature, though some species may modulate incubation initiation and duration in relation to temperature (Coe, Beck, Chin, Jachowski, & Hopkins, 2015; Simmonds, Sheldon, Coulson, & Cole, 2017) and there are reports of GFWO clinging to the sides of the cavity which could be an attempt to reduce heat transfer (Skutch, 1969). Nest temperature is also affected by nest site location and cavity design (although not always) (Butler, Whitman & Dufty, 2009; Zingg, Arlettaz & Schaub, 2010; Sonnenberg, Branch, Benedict, Pitera & Pravosudov, 2020).

Tree decay, in particular, affects thermoregulation of the nest cavity, in that live trees - with higher water content-provide greater insulation against high and low temperature extremes (Grüebler, Widmer, Korner-Nievergelt & Naef-Daenzer, 2014). However, the same trait that makes live trees good insulators also makes them more costly to excavate; on average, live trees are denser than partially dead or decaying trees. Therefore, these birds may be facing an energetic trade-off; whether to put additional effort into excavating a dense live tree-which has higher water content and is better able to thermoregulate eggs and nestlings- or save time and energy by excavating a less stable decayed tree and risk eggs and nestlings overheating.

This possible role for temperature in nest site selection and structure is further strengthened by the trend we observed in vegetation cover, with cavity nesters like the GFWO and the BEWR having higher success in cavities with increased vegetation cover. While the effect size for vegetation (β ranged from 0.02 to 0.05) seems small at first, across the large range of possibilities for cover (1-100) this variable showed a strong effect. For example, with a 15 percent increase in vegetation cover, the effect size for the BEWR grew to 0.30 and the same increase in vegetation cover for the GFWO resulted in an effect size of 0.75, rivaling that of stronger predictors such as decay and cavity cover. Again, these results contrast with previous literature on cavity nesters which indicated a preference for exposed cavities due to increased visibility of approaching predators (Mannan et al., 1980; Li & Martin, 1991; Loye & Carroll, 1998; Newlon, 2005; Jusino et al., 2016). Vegetated cavities in this region may provide increased shade and thus reduced internal temperatures, resulting in another tradeoff, one between temperature regulation and predation.

### Conclusion

Here we evaluated the link between food resources and an ecosystem engineer, and the subsequent influence of this engineer on local secondary cavity nesters. We observed that GFWO nest site location and home range size was positively correlated to biomass of the same three orders of insects that make up large proportions of their diet, and that all SCB had higher nest success in abandoned woodpecker cavities than natural cavities. Thus, GFWO nest in areas with abundant food and SCB reap the benefits of the stable cavities they leave behind, along with opportunistically high insect loads. Our results also suggest that GFWO nest characteristics may influence nest success in ways that differ from more temperate species, indicating future research avenues into energetics and predation pressure tradeoffs in high temperature regions. Additionally, management for woodpeckers and SCB in southern Texas should not focus on the availability of snags (a common management strategy for woodpeckers in temperate climates), but on the number of live trees with a DBH wide enough for nesting.

## Acknowledgments

We thank the East Foundation monitoring teams from 2018 and 2019 for their assistance with data collection. Also, thanks to Andrea Montalvo for her extended field support during data collection. This is manuscript number 056 of the East Foundation.

## Data availability

Data and analytical code are available on Figshare: https://doi.org/10.6084/m9.figshare.12939992.v1 (Hardin et al. 2020)

