## Supplement for "Secondhand homes: Woodpecker cavity location and structure influences secondary nester’s success"

### Supplementary

Variables used throughout manuscript and supplementary materials:

- height = height of the nest measured from the center of the cavity opening to the base of tree
- DBH = diameter at breast height of the tree
- opening = diameter of the cavity opening
- depth = depth of the cavity
- decay = decay ranking, where a rank of one indicated a live tree and rank seven indicated a dead tree with no branches, bark, and soft stem
- vegetation cover = percent of vegetation covering the cavity opening.
- Cavity Type = whether it was in an abandoned woodpecker cavity or a natural cavity.

S. Table 1: Candidate Models for Model Averaging.

| **Bewick's Wren (n = 79)** | logLik | AICc | ΔAICc | Weight |
|---|---|---|---|---|
| M1: Decay + vegetation cover + Cavity Type | -11.672 | 32.520 | 0.000 | 0.185 |
| M2: Decay+ Cavity Type | -13.420 | 33.526 | 1.005 | 0.112 |
| M3: Decay + vegetation cover + Cavity Type + height | -11.465 | 34.748 | 2.227 | 0.061 |
| M4: Decay + vegetation cover + Cavity Type + DBH | -11.498 | 34.815 | 2.295 | 0.059 |
| M5: Decay + vegetation cover+ Cavity Type + opening | -11.573 | 34.964 | 2.443 | 0.054 |
| M6: Decay + vegetation cover + Cavity Type + depth | -11.650 | 35.117 | 2.597 | 0.050 |
| M7: Decay + Cavity Type + DBH | -13.008 | 35.193 | 2.673 | 0.049 |
| M8: Decay + Cavity Type + depth | -13.381 | 35.938 | 3.418 | 0.033 |

| | | | | |
|---|---|---|---|---|
| M9: Decay + Cavity Type + height | -13.412 | 36.001 | 3.481 | 0.032 |
| M10: Decay + Cavity Type + opening | -13.416 | 36.009 | 3.489 | 0.032 |
| M11: Decay + vegetation cover | -15.113 | 36.912 | 4.391 | 0.021 |
| M12: Decay + vegetation cover + Cavity Type + DBH + height | -11.189 | 37.002 | 4.482 | 0.020 |
| *Null Model* | *-47.650* | *97.35* | *31.3106* | *3E-08* |

| **Ash-throated/Brown-crested Flycatcher (n = 102)** | logLik | AICc | ΔAICc | Weight |
|---|---|---|---|---|
| M1: Decay + DBH + Cavity Type + opening | -38.417 | 87.458 | 0.000 | 0.127 |
| M2: Decay + Cavity Type + opening | -39.564 | 87.541 | 0.083 | 0.121 |
| M3: Decay + vegetation cover + Cavity Type + opening | -39.233 | 89.090 | 1.632 | 0.056 |
| M4: Decay + vegetation cover + Cavity Type + DBH + opening | -38.134 | 89.152 | 1.694 | 0.054 |
| M5: Decay + Cavity Type + depth + opening | -39.308 | 89.242 | 1.784 | 0.052 |
| M6: Decay + Cavity Type + DBH + height + opening | -38.252 | 89.387 | 1.929 | 0.048 |
| M7: Decay + Cavity Type + DBH + depth + opening | -38.261 | 89.406 | 1.948 | 0.048 |
| M8: Decay + Cavity Type + DBH | -40.561 | 89.535 | 2.077 | 0.045 |
| M9: Decay + Cavity Type + height + opening | -39.455 | 89.535 | 2.077 | 0.045 |
| M10: Decay + Cavity Type | -41.766 | 89.777 | 2.319 | 0.040 |
| M11: Decay + vegetation cover + Cavity Type + depth + opening | -38.959 | 90.802 | 3.343 | 0.024 |
| M12: Decay + vegetation cover + Cavity Type + height + opening | -39.023 | 90.930 | 3.472 | 0.022 |
| M13: Decay + vegetation cover + Cavity Type + DBH + height + opening | -37.872 | 90.936 | 3.477 | 0.022 |
| M14: Decay + vegetation cover + Cavity Type + DBH + depth + opening | -37.957 | 91.106 | 3.647 | 0.020 |
| M15: Decay + Cavity Type + DBH + height | -40.277 | 91.178 | 3.720 | 0.020 |
| M16: Decay + Cavity Type + depth + height + opening | -39.162 | 91.209 | 3.751 | 0.019 |
| M17: Decay + Cavity Type + DBH + depth + height + opening | -38.060 | 91.312 | 3.854 | 0.018 |
| M18: Decay + vegetation cover + Cavity Type + DBH | -40.344 | 91.313 | 3.854 | 0.018 |

| | | | | |
|---|---|---|---|---|
| M19: Decay + vegetation cover + Cavity Type | -41.503 | 91.419 | 3.961 | 0.017 |
| M20: Decay + Cavity Type + height | -41.549 | 91.511 | 4.052 | 0.017 |
| M21: Cavity Type + opening | -42.719 | 91.682 | 4.224 | 0.015 |
| M22: Decay + Cavity Type + DBH + depth | -40.557 | 91.738 | 4.280 | 0.015 |
| M23: Decay + Cavity Type + depth | -41.751 | 91.914 | 4.456 | 0.014 |
| *Null Model* | *-67.350* | *136.74* | *49.2824* | *2.5E-12* |

| **Golden-fronted Woodpecker (n = 55)** | logLik | AICc | ΔAICc | Weight |
|---|---|---|---|---|
| M1: Decay + vegetation cover | -12.655 | 31.781 | 0.000 | 0.267 |
| M2: Decay + vegetation cover + DBH | -12.319 | 33.439 | 1.657 | 0.117 |
| M3: Decay + vegetation cover + opening | -12.607 | 34.014 | 2.233 | 0.087 |
| M4: Decay + vegetation cover + height | -12.636 | 34.072 | 2.290 | 0.085 |
| M5: Decay + vegetation cover + depth | -12.640 | 34.080 | 2.299 | 0.085 |
| M6: Decay + vegetation cover + DBH + depth | -12.207 | 35.639 | 3.857 | 0.039 |
| M7: Decay + vegetation cover + DBH + opening | -12.249 | 35.723 | 3.941 | 0.037 |
| M8: Decay + vegetation cover + DBH + height | -12.319 | 35.862 | 4.081 | 0.035 |
| M9: Decay + vegetation cover + height + opening | -12.567 | 36.358 | 4.577 | 0.027 |
| *Null Model* | *-32.227* | *66.530* | *34.749* | *7.6E-09* |

| **Black-crested Titmouse (n = 39)** | logLik | AICc | ΔAICc | Weight |
|---|---|---|---|---|
| M1: Decay + vegetation cover + Cavity Type | -11.672 | 32.520 | 0.000 | 0.185 |
| M2: Decay + Cavity Type | -13.420 | 33.526 | 1.005 | 0.112 |
| M3: Decay + vegetation cover + Cavity Type + height | -11.465 | 34.748 | 2.227 | 0.061 |
| M4: Decay + vegetation cover + Cavity Type + DBH | -11.498 | 34.815 | 2.295 | 0.059 |
| M5: Decay + vegetation cover + Cavity Type + opening | -11.573 | 34.964 | 2.443 | 0.054 |
| M6: Decay + vegetation cover + Cavity Type + depth | -11.650 | 35.117 | 2.597 | 0.050 |
| M7: Decay + Cavity Type + DBH | -13.008 | 35.193 | 2.673 | 0.049 |
| M8: Decay + Cavity Type + depth | -13.381 | 35.938 | 3.418 | 0.033 |
| M9: Decay + Cavity Type + height | -13.412 | 36.001 | 3.481 | 0.032 |

| | | | | |
|---|---|---|---|---|
| M10: Decay + Cavity Type + opening | -13.416 | 36.009 | 3.489 | 0.032 |
| M11: Decay + vegetation cover | -15.113 | 36.912 | 4.391 | 0.021 |
| M12: Decay + vegetation cover + Cavity Type + DBH + height | -11.189 | 37.002 | 4.482 | 0.020 |
| *Null Model* | *-26.917* | *55.940* | *23.422* | *1.52E-06* |

#### Supp Figs.

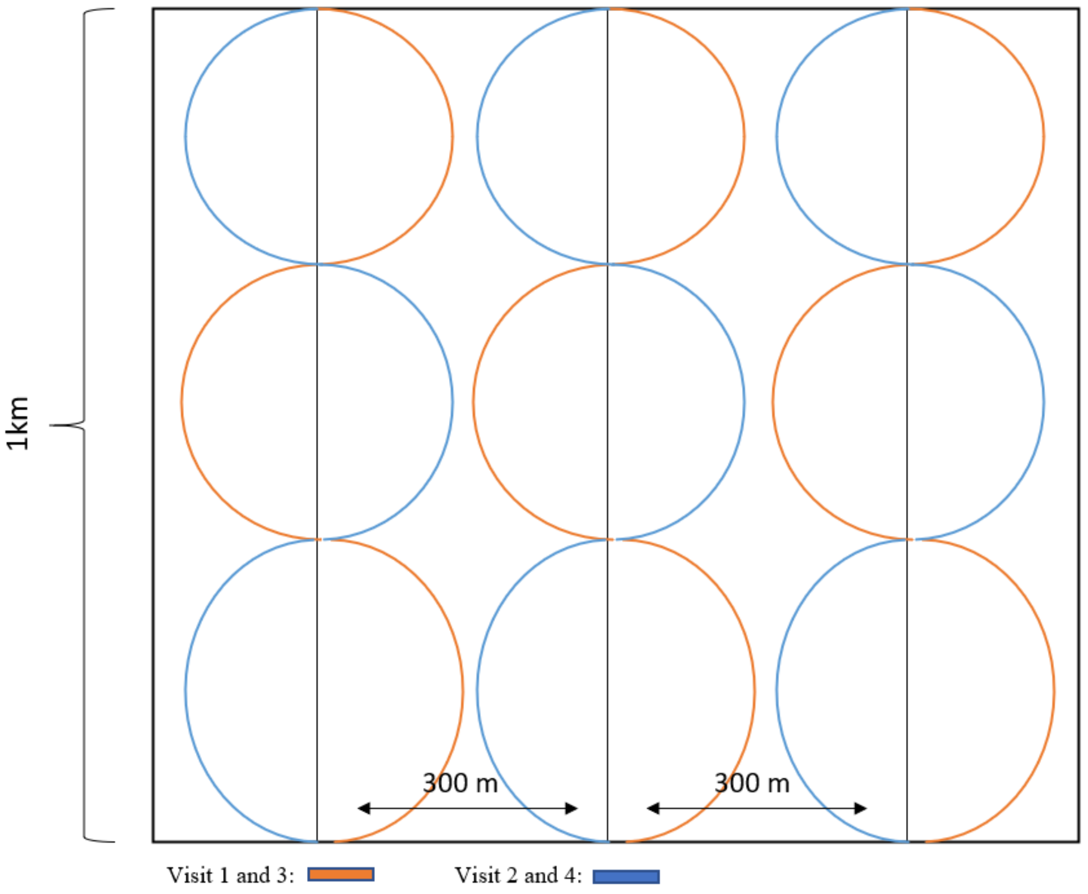


**Sup Fig. 1: Visual of walking transects used to detect and locate Golden-fronted woodpecker nests throughout the 12, 1 km plots placed on the San Antonio Viejo Ranch, East Foundation, during spring and summer 2019. Orange path was followed during visits 1 and 3, blue path was followed for visits 2 and 4.**

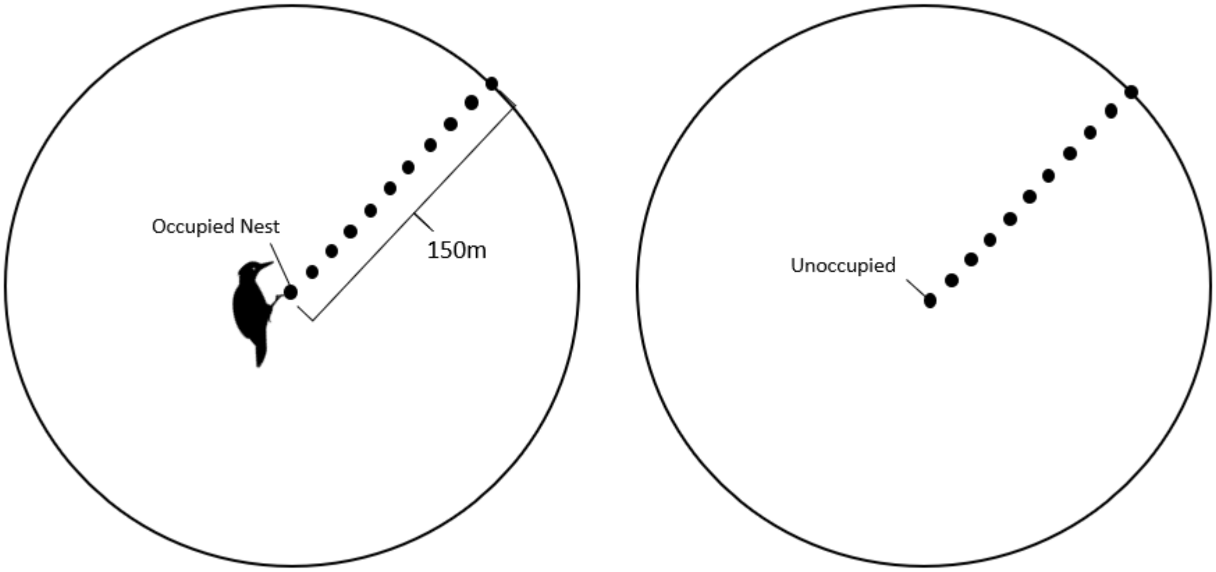


**Supp Fig 2: Insect sampling method: A subset of 24 Golden-fronted woodpecker nests (occupied), and their corresponding unoccupied sites (having same vegetation alliance) was chosen to conduct insect surveys. Insects were sampled once a week from May-July with a sweep net, samples were sorted to insect order, and dried and weighted.**
